# Probing EEG Foundation Model Attention Maps for Lightweight Interpretable Emotion Recognition

**DOI:** 10.64898/2026.11.05.764629

**Authors:** L. Arvanitopoulos, A. Dragomir, C. Fidas, A. Bezerianos

## Abstract

Foundation Models (FMs) trained on EEG signals promise subject- and dataset-independent pipelines for brain-computer interface (BCI) applications, but have yet to consistently outperform task-specific architectures, and their large representations are costly to compute and store. As a means of advancing the understanding of the Foundation Models’ inner mechanisms, we propose a lightweight alternative that exploits the self-attention mechanism of REVE, a pretrained EEG FM, to derive features similar to weighted Functional Connectivity (FC) without any fine-tuning. Inspired by kernel-based FC matrices, we aggregate the raw first-layer attention weights of a single head over the full duration of dataset trials into weighted channel-to-channel adjacency matrices. These matrices replace standard encoder representation vectors as classification features, substantially reducing computational cost. We further introduce a label-free, symmetry-based criterion, motivated by the emergence of near-symmetric attention patterns in trained encoder-decoder transformers, ranking attention heads by how much discriminative information they carry. Classical machine learning classifiers are then trained on the single most informative head’s attention-derived FC matrix and achieve balanced accuracy and F1 scores on the SEED-IV emotion-recognition dataset that are competitive with several fully fine-tuned state-of-the-art EEG Foundation Models, requiring at least two orders of magnitude fewer calculations than the standard use of the encoder. Subsequent cortical region analysis revealed a significant over-representation of frontal region connections by ranking the most discriminative FC matrix connectivity edges, validating our methods as consistent with prior affective-computing literature. These results suggest that interpretable, computationally inexpensive linear probing of attention maps is a viable route to deploying EEG Foundation Models in resource-constrained settings.

## 1 Introduction

Recurrence-free attention based models [1], led by transformers, have consistently been the state of the art (SOTA) in multiple tasks, making Foundation Models (FMs) effectively possible, either in the form of Large Language Models (LLMs) in Natural Language Processing (NLP) or in other fields, such as neuroscience [2]. Some investigation of the attention mechanism and emergent behavior has been done on LLMs, with the goal of understanding the supremacy of the transformer architecture over others in the field of NLP. Replication of those studies in other fields is even more scarce, with more literature focusing on the outputs of the models, rather than the inner mechanisms. In the analysis of neurophysiological data, FMs attempt to mitigate inter-subject and cross-experimental variability of neurophysiological experiment pipelines [3]. Despite their success in other modalities, FMs trained on brain signals have yet to out-perform task-specific or time-series FM models [4]. Understanding the inner workings of the FMs can help in optimizing their use on neuroscience trial pipelines, by reducing redundant processing that only adds further noise and computational cost.

The Human-Computer-Interaction (HCI) research’s long sought goals are to operate machines with minimal effort, or to objectively characterize human state. Brain-Computer-Interfaces (BCIs) were developed as such, to capture brain activity and turn it into actionable measures, but also take it a step further by attempting to use subconscious states as control signals. Affective computing extends this effort to human emotion, incorporating it into the pipeline process. Finally, the biggest stepping stone in understanding the affective state of the user, is the task of Emotion Recognition, the solution of which would provide a complete, bidirectional communication between man and machine.

In this context, electroencephalography (EEG) is the most prominent technique for recording brain activity, due to its portability and cost-effectiveness. EEG’s ease of use and its excellent temporal resolution makes it an ideal technique for interactivity and rapid data collection, a prerequisite to the concept of BCI [5]. However, the wide-spread adoption of EEG remains challenging, particularly due to the high individual variability in the data, as well as issues such as relatively low signal to noise ratio, or reproducibility across recording sessions [6]. In this sense, neuroscience-focused FMs hold the promise of better accounting for these issues. Typically, these models are pre-trained on public EEG datasets consisting of multiple BCI tasks, often differing in structure or experimental design, due to the large disparity between the amount of data available from in-house experiments and the large amounts of data required for training [7]. Overall, the goal of FMs is to improve the generalizability of cross-subject BCI pipelines, by replacing the classically extracted features with deep learning representations. Several FMs have become a staple in benchmarking contexts [8, 9], with models such as LaBraM [10], EEGPT [11] and NeuroGPT [12] being trained specifically on EEG signals. CSBrain, CBraMod [13, 14] focus on better positional encoding and newer models [15, 16] improving the tokenizers and loss functions. While benchmarking has been the predominant use case, exceptions [3] prove that internalization of FMs in a BCI pipeline can reduce the preprocessing inference time compared to prior typical implementations. In this direction, among the most recent literature of EEG FMs, the REVE model [16] seems to offer the most flexibility with its novel 4D positional encoder and promises optimal linear probing performance. Good linear probing performance constitutes a necessary condition to allow for the realization of pipelines with FMs on environments with less computational power and without hardware acceleration.

The key component of the transformer architecture in FMs is the Multi-Headed Attention (MHA) layer with its self attention mechanism, that enables the models to attend to information from different subspaces and scales [1]. Interpretation of the attention weights, a necessary step of the attention mechanism, as a normalized kernel [17] shows that the MHA layer maps a soft adjacency matrix on the input tokens. In transformer models trained on EEG data [16], the tokens fed in the model can be traced back to a unique channel and time window from the input data. Thus, kernel-based generation of functional connectivity matrices has been proposed as a method of extracting non-linear connections [18], and combined with the inherent symmetry over the diagonal in the encoder-decoder architectures that emerges from training [19], means that attention weights generated in the MHA layer can be converted into functional connectivity matrices. Consequently, extraction of channel to channel connections from the attention weights maps can lead to uncovering features similar to soft functional connectivity and is the focus of our present work.

Functional connectivity analysis has been applied as feature extraction for cross-subject emotion recognition [20], typically proving to yield more robust sets of features. However, functional connectivity is a relatively complex procedure, requiring extensive processing steps, some of which with outcomes that depend on experimenters’ choices (e.g. estimator methods, network density thresholding etc). Further, the robustness of functional connectivity features is typically affected by the level of noise present in the data.

Explanation of the superiority of the transformer architecture has been attempted in literature, either by tracing the internal calculations to previous machine learning concepts or by examining emerging behavior. Specifically, one approach is performed through mathematical analysis of the internal mechanisms, linking it to classical methods such as kernels [17], whereas others detect emerging patterns of clustering and symmetry [21] [19] in the attention maps of pretrained models. However, similar attempts in explaining neuroscience-based FMs internal mechanisms and retrieved representations have so far been limited [22]. These observations motivate the contributions of this paper.

In this paper we apply the features derived from attention maps to classical methods of machine learning classifiers and demonstrate performance on par with other implementations of Foundation Models for the SEED-IV dataset [23], requiring only a fraction of the usual architecture. To achieve this, we also propose the use of the symmetry scores on specific transformations of each head, to rank the discriminative ability of each attention map in the context of emotion recognition.

## 2 Methods

To minimize out-of-distribution and domain-shift errors that can occur from datasets outside the FM’s pretraining data, we use the SEED-IV dataset from the SJTU Emotion EEG Dataset (SEED) collection [24]. Each dataset in the SEED collection is built from EEG recordings of subjects stimulated with film clips designed to elicit specific emotions, which are subsequently used as the label of each corresponding sample. SEED-IV contains four labels, ‘Neutral’, ‘Sad’, ‘Fear’ and ‘Happy’. The EEG recordings were acquired with the 62-channel ESI NeuroScan System over three sessions, each consisting of 24 trials, for each of 15 subjects. This yields 3 *×* 24 = 72 trials per subject and 72 *×* 15 = 1,080 samples in total [24]. Because the film clips vary in duration, the dataset is naturally represented as a tensor of Samples *×* Channels *×* Timepoints, with a variable number of timepoints per sample.

We applied bandpass filtering in the range 0.5-100 Hz and the signals were resampled to 200 Hz (down from the original 1kHz) to match the REVE model’s input requirements [16]. Examination of the frequency content of the recordings indicated no measurable powerline interference, so no notch filtering was required. To preserve fairness with respect to other FMs benchmarked on the same dataset, no additional artifact rejection was performed. Finally, the CB1 and CB2 electrodes were dropped, since they fall outside REVE’s positional bank, leaving 60 usable channels per sample. After processing, each recording is split into *W* windows of 1 second, as part of the model’s tokenization procedure.

### 2.1 Attention Scores as Kernels and their Interpretability

Among recent EEG Foundation Models, REVE [16] was selected because it promises improved linear-probing performance, which is a necessary condition for deploying FM-based pipelines in environments without hardware acceleration. REVE adopts an encoder-decoder transformer architecture together with a novel four-dimensional positional encoding stage that allows the model to generalize to electrode montages that were not necessarily seen during pretraining. Because REVE is transformer-based, it relies on the self-attention mechanism to compute soft connections between the input tokens produced from spatiotemporal windows of the EEG signal.

Attention scores can be interpreted as a kernel (1) over the input tokens [17]: for query and key projection matrices *W*_*Q*_ and *W*_*K*_ of tokens *x*_*i*_ and *x*_*j*_.

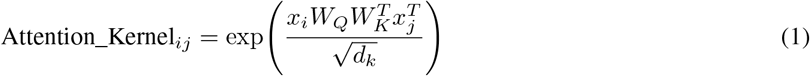

The resulting attention weights are simply this kernel normalized, denoted in literature by the softmax function of the attention scores, which maps values onto [0, 1]. Because tokens correspond to channel/window embeddings of the input recording, aggregating attention weights across the temporal dimension converts a per-head attention map into an attention-derived adjacency matrix over channels. Similarities between signals recorded at different brain regions are conventionally interpreted in neurophysiological analysis as soft connections akin to functional connectivity. Building on recent evidence that near-diagonal symmetry emerges in the attention weights of trained encoder-decoder transformers [19], and given that functional connectivity analysis itself can be defined through a similarity metric such as a kernel [18], we treat the attention weights of REVE’s early layers as candidate functional connectivity weights.

It must be noted that this interpretation is restricted to the first attention layer of the encoder. The MHA layer in a standard transformer block is followed by a linear (output) projection [1] that mixes information across heads and positions; from the second layer onward, this mixing means that an individual attention score can no longer be unambiguously attributed to a single channel/window pair. Only in the first layer does each token still correspond exactly to one channel-window sample of the input, which is what allows the resulting attention maps to be read as interpretable, channel-resolved connectivity matrices.

### 2.2 From Multi-Head Attention to Soft Functional Connectivity

Starting from the 1,080 SEED-IV recordings (3 sessions *×* 24 trials *×* 15 subjects) of size 60 channels *×W* windows, the channel names of the remaining 60 electrodes are passed through REVE’s positional bank to obtain their positional encodings. Each sample is then forward-passed through REVE without any fine-tuning, with hooks embedded in the PyTorch model to capture the raw *QK*^*T*^ values produced by every head of the first attention layer.

Figure 1 presents an overview of the proposed pipeline, with the model being used for the extraction of soft functional connectivity weights that are then fed into classical ML classifiers. The raw *QK*^*T*^ attention scores are passed through the softmax function to obtain attention weights, which reduces their dynamic range to a value comparable to that of typical functional connectivity metrics. For a trial with *W* windows, the resulting attention maps have shape 8 heads *×*60*W ×*60*W*, since each token multiplexes a channel and a window. The first step is to exclude the diagonal channel-channel regions, since they infer self connections. To collapse the matrix into a single soft adjacency matrix per head, we max-pool the attention map with a stride equal to the number of windows in the trial, aggregating over the entire trial length. This produces, per trial and per head, a 60*×*60 attention-generated soft functional connectivity matrix, and eight such matrices per trial in total (one per attention head). The visualization in Figure 2, displays the required steps from the extraction of the softmax values into a weighted soft functional connectivity matrix. Specifically, the first stage is extracting the raw *QK*^*T*^ scores, normalized with the softmax function, i.e. the attention weights of channel-window tokens (Fig. 2A), followed by zeroing out self connections of channels (squares on the diagonal) (Fig.2B). Then averaging the individual attention weights over the time windows within each trial leads to the aggregated soft connectivity matrices shown in Fig 2C and D, each square holding the connection weight between 2 channels aggregated throughout the trial. The aggregated matrix can then be converted to a functional connectivity-like graph through sparsity thresholding, to facilitate results visualization (Fig. 2E) The first three plots (Fig. 2A, B, and C) are magnified to show only the first 8 EEG channels for clarification purposes.

**Figure 1.**
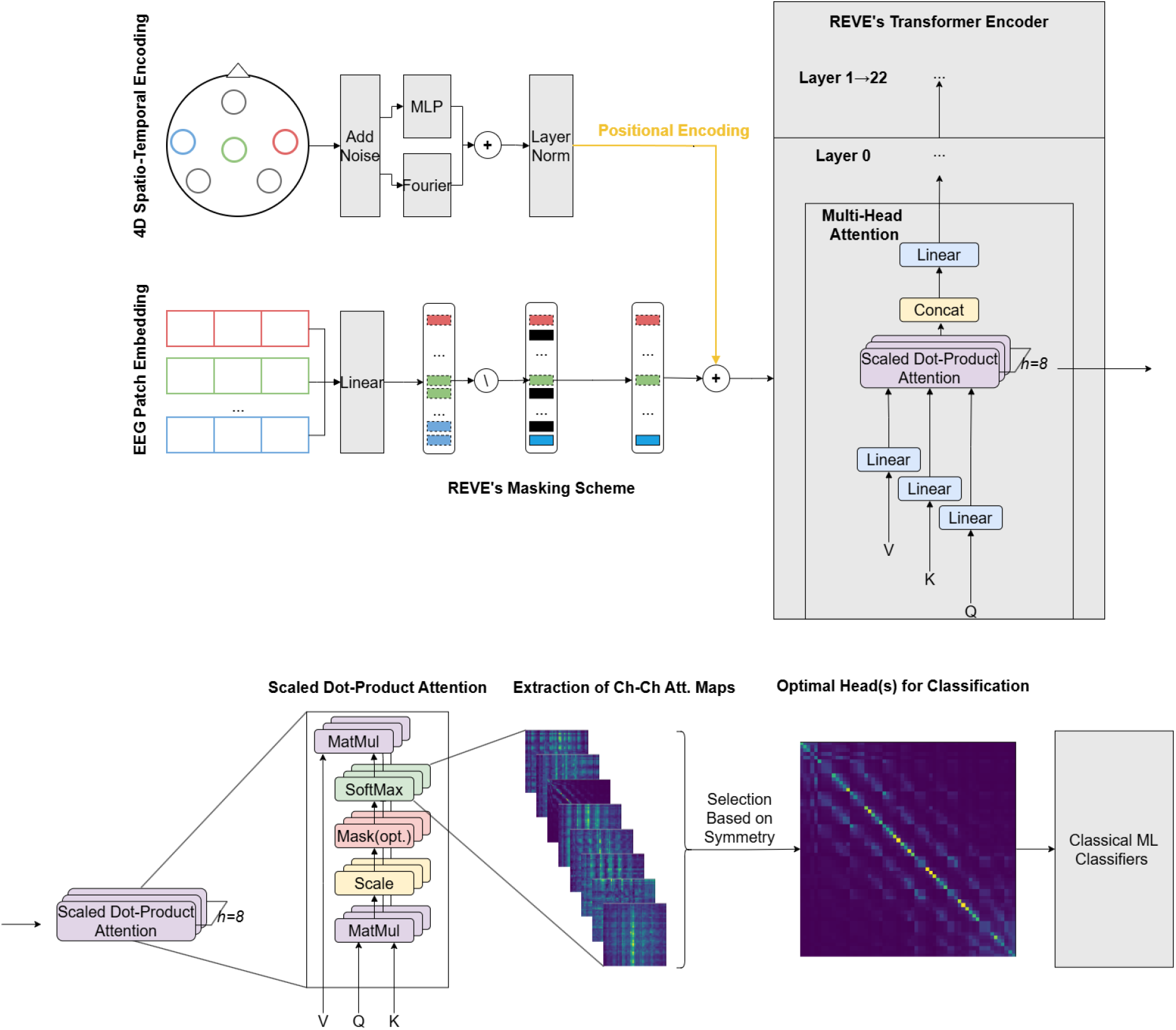
1^*st*^ Overview of the proposed pipeline: extraction of attention maps from the first layer of the encoder stage as features for subsequent analysis

**Figure 2.**
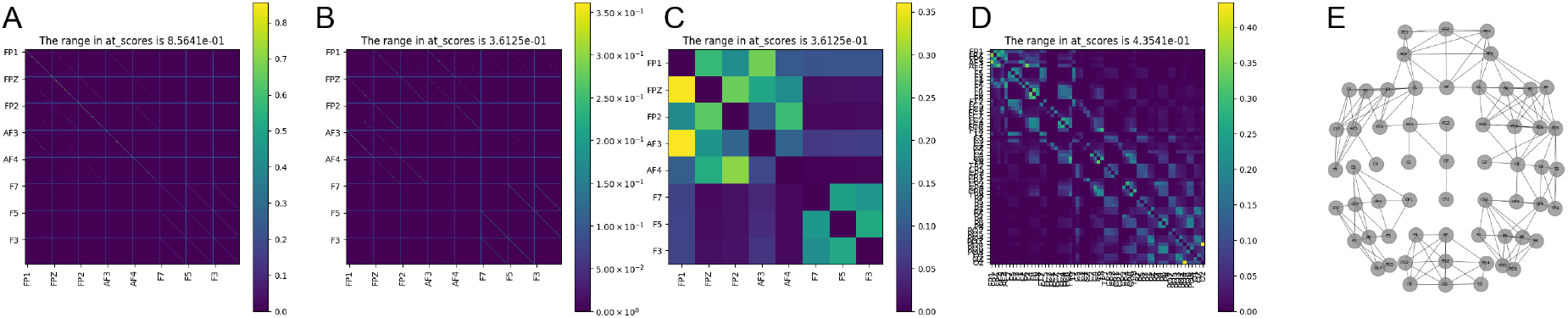
The first three plots (A, B and C) are magnified plots corresponding to the first 8 channels of a specific attention head, while the fourth plot (D) displays the full matrix corresponding to the plot in C. This matrix can then be converted to a functional connectivity-like graph through sparsity thresholding (E).

### 2.3 Quantifying Discriminative Information in Attention Heads via a Symmetry Scores

To identify which of the eight attention heads carries the most actionable information, and should therefore be used for downstream analyses, we apply sparsity thresholding to each head’s 60*×*60 matrix, retaining only the strongest *p*% of the 3,600 candidate connections, and binarize the retained connections to 1. We then define a symmetry score and a symmetry error by applying, for every binarized map, along the main diagonal (2) the Frobenius norm in the difference of each matrix with respect to its own transpose.

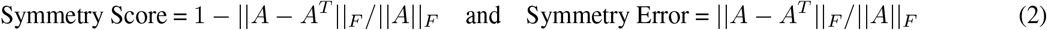

The definition of quantitive measure such as symmetry score and error, allows objective comparisons of our observations of diagonal symmetry through their values. Selection of measures exploiting matrix norms was the obvious choice for reducing the nature of the matrix and its transpose into scalar values.

### 2.4 Classification and Evaluation

After extracting the attention-generated weighted soft functional connectivity matrices, we flatten each trial’s 60*×*60 matrix, giving a feature matrix of 1,080 samples *×*3,600 features per head. Subsequently, to benchmark this representation against the standard way of extracting features from REVE, we also compute two attention-pooling baselines. The first follows REVE’s recommended feature-extraction procedure [16], concatenating the attention-pooled embeddings of all 23 intermediate layer outputs, producing a 23*×*512 = 11,776-dimensional feature vector per trial. The second uses only the final layer’s pooled representation, giving a much smaller 512-dimensional feature vector per trial.

For the classification stage, a panel of classical machine learning classifiers are trained in parallel on each of the feature sets above (i.e., the attention-pooled embeddings, the final layer’s representation and the soft functional connectivity features obtained following our methodology), with their results compared in Section 3. These classifiers include linear approaches (logistic regression, linear-kernel SVM), an RBF-kernel SVM, boosting algorithms (XGBoost, LightGBM), an ensemble method (Random Forest), and simple neural networks. All classifiers are evaluated under a subject-independent Leave-One-Subject-Out (LOSO) protocol, and compared using accuracy, area under the curve (AUC), F1-score and Hamming loss, among other standard metrics. As SEED-IV is a balanced four-class problem (*K* = 4), a random classifier that predicts each class with equal probability achieves an expected accuracy, recall and precision of 1*/K* = 0.25, an expected F1-score of 0.25, and an expected Hamming loss of 1*−* 0.25 = 0.75; we use these values as the chance-level baseline throughout.

## 3 Results

### 3.1 Symmetry as Metric of Attention Head Importance

We first evaluate the symmetry criterion used to rank attention heads, according to their discriminative value. Across heads, we observe a clear progression toward more distinct, more diagonally-symmetric maps, and identify the third head (index 2) as carrying the most information under the symmetry-based criterion. We validate this ranking downstream by contrasting the classification performance obtained from the third head’s features against that of the remaining seven heads. Figure 3 reports the statistical momentums of the Frobenius symmetry error and score, computed on the sparsity-thresholded binarized first-layer attention maps with value of 10% and aggregated across all trials, for each of the eight heads. From left to right, the values are processed before applying symmetry scores. In 3A there is no processing, B is after zeroing anything below the 10% strongest values and finally, C is with binarizing the values of the matrices to 1. The third head (index 2) attains the lowest mean Frobenius symmetry error and the highest mean symmetry score, together with the lowest standard deviation on both metrics, indicating that it produces the most consistently symmetric, and hence most functional connectivity-like, attention pattern.

**Figure 3.**
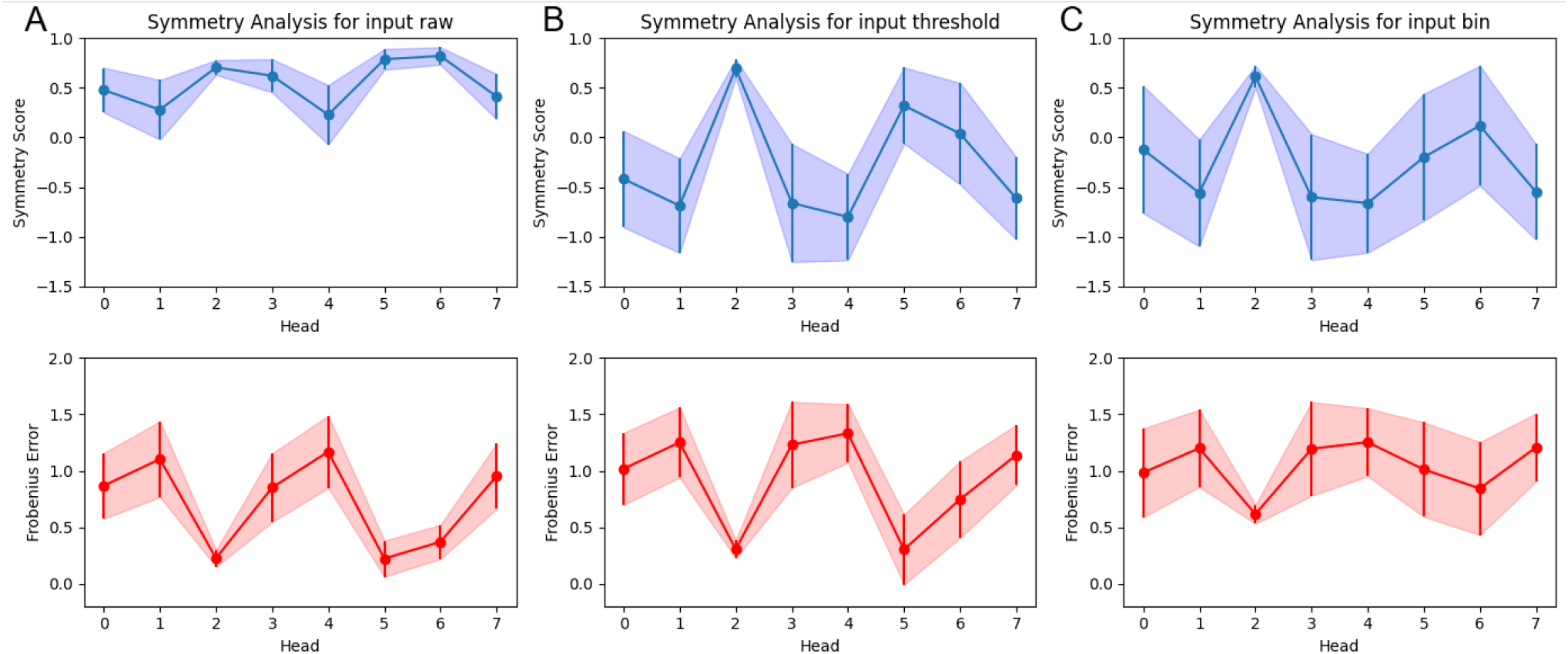
Frobenius symmetry error and score (mean and standard deviation) computed on the raw first-layer attention maps of each of the eight heads, aggregated over all trials, then on zeroed elements after sparsity-thresholding and finally binarized version of the latter.

To qualitatively visualize this difference, Figure 4 shows examples of attention-generated maps for heads 0, 2 and 4. The map produced by head 2 (Fig. 4B) exhibits a visibly more block-structured and diagonally symmetric pattern than heads 0 or 4, consistent with the quantitative symmetry scores above.

**Figure 4.**
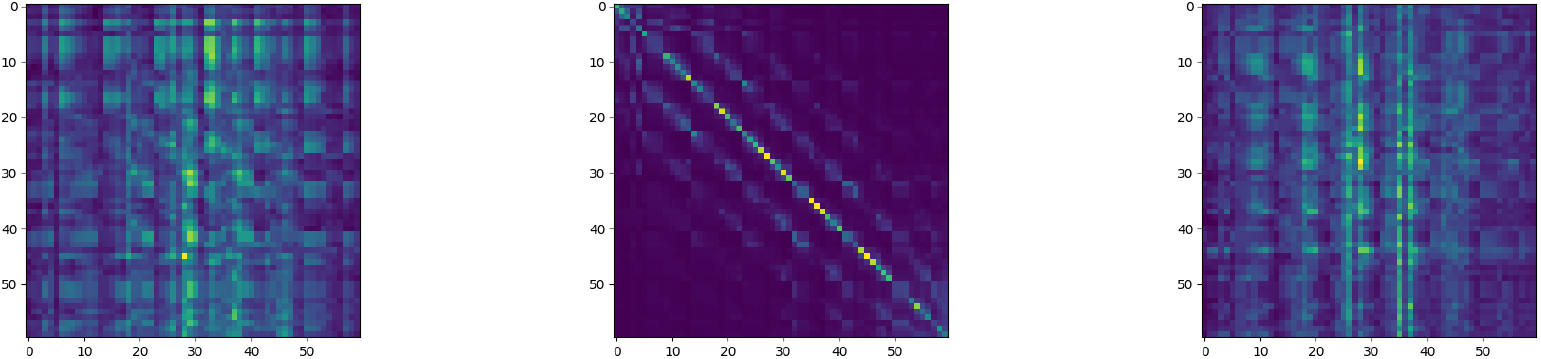
Attention-generated Functional Connectivity maps for heads 0, 2 and 4 of the first attention layer.

Table 1 summarizes accuracy, precision, recall, F1-score and Hamming loss for all eight heads individually, together with the chance-level baseline. Head 2 achieves the best performance across every metric (accuracy 0.3917, F1 0.3803), followed by head 4 as the second-best head, while the remaining heads cluster closer to, though still consistently above, the 0.25 chance baseline. Permutation testing the features of head 2 allows us to reject the null hypothesis that random features can perform similarly with p<0.01. This confirms that the symmetry-based ranking identified in Figure 3, obtained without using any labels, correctly identifies the most discriminative attention head for the downstream emotion-recognition task.

**Table 1:** Performance comparison of the eight attention heads on the SEED-IV dataset.

| Head | **Accuracy ↑** | **Precision ↑** | **Recall ↑** | **F1 Score ↑** | **Hamming Loss ↓** |
|---|---|---|---|---|---|
| 0 | 0.3250 | 0.3551 | 0.3250 | 0.3039 | 0.6750 |
| 1 | 0.2917 | 0.2670 | 0.2917 | 0.2372 | 0.7083 |
| 2 | **0.3917** | **0.3863** | **0.3917** | **0.3803** | **0.6083** |
| 3 | 0.3278 | 0.3371 | 0.3278 | 0.2833 | 0.6722 |
| 4 | <u>0.3639</u> | <u>0.3787</u> | <u>0.3639</u> | <u>0.3532</u> | <u>0.6361</u> |
| 5 | 0.2667 | 0.2266 | 0.2667 | 0.2323 | 0.7333 |
| 6 | 0.3056 | 0.2084 | 0.3056 | 0.2061 | 0.6944 |
| 7 | 0.3167 | 0.3017 | 0.3167 | 0.2672 | 0.6833 |
| Baseline | 0.2500 | 0.2500 | 0.2500 | 0.2500 | 0.7500 |

### 3.2 Comparison with Other Methods

Given that SEED-IV is a balanced four-class classification problem (*K* = 4), the baseline performance in Table 1 corresponds to a random classifier that predicts each class with equal probability. In Table 2 we compare the metric heatmaps of the attention-generated weighted FC features from head 2 against the two attention-pooling baselines described in Section 3 (i.e., the attention-pooled embeddings, the final layer’s representation and the soft functional connectivity features obtained following our methodology). Thus, we can contrast our features with attention pooling concatenated across all 23 layers (11,776 dimensions) and with attention pooling from the last layer (only 512 dimensions), across the same set of classifiers and metrics.

**Table 2:** Performance Metrics of All Model Configurations and Features, Color-Graded from Worst (red) to Best (green)

| **Classifier Model** | **Input Features** |  |  | **Attention Pools (All Layers)** |  |  | **Attention Pools (Final)** |  |  |
|---|---|---|---|---|---|---|---|---|---|
|  | **Acc** | **F1** | **AUC** | **Acc** | **F1** | **AUC** | **Acc** | **F1** | **AUC** |
| Logistic Regression | 0.376 | 0.348 | 0.672 | 0.342 | 0.332 | 0.597 | 0.347 | 0.335 | 0.618 |
| SVM (Linear) | 0.374 | 0.345 | 0.676 | 0.346 | 0.336 | 0.506 | 0.342 | 0.326 | 0.511 |
| SVM (RBF) | 0.393 | 0.346 | 0.664 | 0.254 | 0.110 | 0.469 | 0.248 | 0.103 | 0.379 |
| Random Forest | **0.421** | **0.401** | 0.696 | 0.356 | 0.335 | 0.621 | 0.340 | 0.312 | 0.618 |
| Gradient Boosting | 0.381 | 0.361 | 0.669 | 0.355 | 0.332 | 0.627 | 0.346 | 0.327 | 0.634 |
| XGBoost | 0.394 | 0.364 | 0.693 | 0.350 | 0.333 | 0.640 | 0.347 | 0.330 | 0.625 |
| LightGBM | 0.410 | 0.386 | **0.698** | 0.381 | 0.358 | 0.656 | 0.359 | 0.337 | 0.630 |
| AdaBoost | 0.369 | 0.347 | 0.671 | 0.304 | 0.285 | 0.606 | 0.330 | 0.310 | 0.606 |
| KNN (*k* = 5) | 0.299 | 0.254 | 0.560 | 0.287 | 0.261 | 0.531 | 0.296 | 0.282 | 0.558 |
| Neural Net. (128) | 0.405 | 0.360 | 0.683 | 0.266 | 0.259 | 0.509 | 0.248 | 0.213 | 0.508 |
| Neural Net. (256-128) | 0.388 | 0.335 | 0.676 | 0.269 | 0.256 | 0.514 | 0.267 | 0.234 | 0.534 |

In both comparisons, attention-generated weighted soft functional connectivity features from head 2 match or exceed the performance obtained with the much higher-dimensional attention-pooling baselines, despite requiring 176*×* and 8*×* fewer calculations than the all-layer and last-layer pooling strategies, respectively. The features were preprocessed with standard scaling and PCA dimensionality reduction down to 200 components to improve compute time.

Finally, we place our method in the context of previously published benchmarks on SEED-IV [8]. Table 3 reports balanced accuracy (B-Acc) and weighted F1-score (F1-W) for a set of traditional and Foundation Model baselines under the cross-subject setting, alongside our method. Due to the SEED-IV dataset being balanced, we can safely compare the metrics between the two studies.

**Table 3:** Benchmarking results in the cross-subject setting. The best and second-best results are bold and underlined, respectively. Baselines other than our method use full fine-tuning; our method uses only linear probing.

| **Metrics** | **Traditional Models** |  |  |  | **EEG Foundation Models** |  |  |  |  |
|---|---|---|---|---|---|---|---|---|---|
|  | **EEGNet** | **LDMA** | **ST-Tran** | **Conformer** | **BIOT** | **EEGPT** | **LaBraM** | **CBraMod** | **Our Method** |
| B-Acc | 34.85 | 36.32 | 32.94 | 34.94 | 35.06 | 31.20 | <u>40.98</u> | 39.36 | **42.13** |
| F1-W | 28.72 | 35.45 | 33.20 | 33.20 | 33.52 | 29.94 | **40.61** | 38.70 | <u>40.12</u> |

Our method achieves the best balanced accuracy (42.13) among all compared models, and the second-best weighted F1-score (40.12), trailing only the fully fine-tuned LaBraM model (40.61). Notably, every other Foundation Model baseline in Table 3 is reported under full fine-tuning, whereas our method uses only linear probing of a frozen backbone together with a single attention head’s FC features, at a fraction of the computational cost.

### 3.3 Frontal Region Over-Representation

To investigate the spatial organization of the discriminative soft functional connectivity patterns identified by head 2, we selected the most discriminative edges using an ANOVA F-score criterion, taking the top 10% strongest (360 edges in total) of all candidate connections. Figure 5 is the sparsity thresholded adjacency matrix (left) and soft connection matrix (right) of the respective edges. We grouped these edges into three categories: connections within the frontal, anterior frontal, frontal, frontotemporal and frontocentral regions (FP/AF/F/FT/FC), connections between these regions and the rest of the scalp, and connections entirely within the remaining regions.

**Figure 5.**
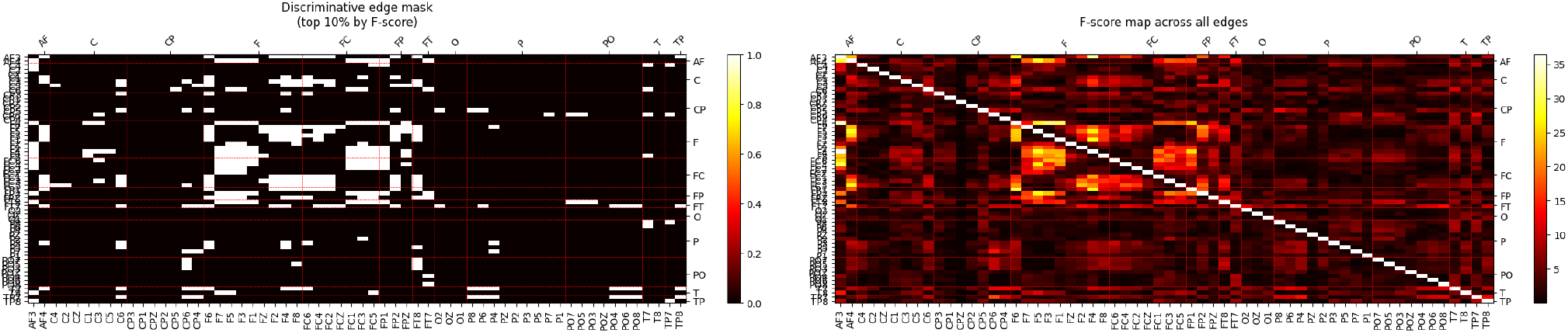
Scalp topography of the most discriminative attention-generated soft functional connectivity edges, showing over-representation of frontal-region electrodes.

As shown in Table 4, edges within the FP/AF/F/FT/FC regions are substantially over-represented, with 215 observed edges against 51.5 expected edges under a uniform distribution, corresponding to a 4.18× enrichment (one-sided hypergeometric test, p < 10^*−*16^). Because edges sharing nodes and neighbouring channels are not independent, we additionally assessed significance with a label-permutation test (n = 5,000), in which the F-score ranking and top-10% selection were repeated on label-shuffled data; the observed count exceeded all permuted counts (p < 0.005). This over-representation of frontal-lobe electrodes among the most discriminative edges is consistent with prior studies on affective computing [25], and suggests that the connectivity patterns learned by REVE’s third attention head are concentrated within frontal-related regions that are already known to be dominant contributors to emotion-recognition performance.

**Table 4:** Regional distribution of the top 10% most discriminative functional connectivity edges. The expected number of edges is estimated according to the overall distribution of candidate edges. The ratio represents the enrichment factor between the observed and expected edge counts.

| **Edge Category** | **Observed** | **Expected** | **Ratio** |
|---|---|---|---|
| Inner FP/AF/F/FT/FC region connections | 215 | 51.5 | 4.18× |
| Between FP/AF/F/FT/FC and other regions | 91 | 173.1 | 0.53× |
| Within other regions connections | 54 | 135.5 | 0.40× |

## 4 Discussion

In the present work we highlight four main contributions. First, we show that raw first-layer attention weights of a pretrained EEG Foundation Model can be repurposed, without any fine-tuning, into explainable classification features similar to a functional connectivity map in its own right. Second, by operating on a single attention head’s 60 *×* 60 matrix rather than pooled embeddings across all layers, the method reduces memory demand by 176 times and computational cost by 22 times at inference relative to standard attention pooling, by bypassing the majority of the model. Third, we propose the use of a label-free, symmetry-based criterion for head selection, which we validate against downstream classification performance. Fourth, by grounding the analysis in classical functional connectivity analysis, the method offers a bridge between the internal mechanisms of EEG Foundation Models and established neurophysiological interpretations of affective processing.

The head-selection results, in particular, offer interesting insight into how discriminative information is distributed across heads. Non-symmetric heads yield close-to-random classification performance and, on inspection, tend to collapse onto only one or two predicted labels, consistent with the poor diagonal structure visible in their attention maps. Head 2 is the only head that develops a more pronounced diagonal structure in comparison to the rest. This correspondence supports the hypothesis that diagonal symmetry in attention weights is a meaningful proxy for the presence of usable, channel-resolved structure, rather than an artifact of the aggregation procedure.

Compared against fine-tuned implementations of other Foundation Models, our method reduces the associated computational cost of the extracted features by a factor of roughly 22 layers *×*8 heads = 176 relative to standard all-layer attention pooling, while also slightly improving classification performance on the SEED-IV dataset. This is a notable result given that every other Foundation Model baseline we compare against relies on full fine-tuning, whereas our approach uses only linear probing of a frozen backbone.

As noted in Section 3, the MHA layer of the transformer architecture is followed by a linear output projection [1]. Interpretability of individual attention scores as channel-to-channel connections is therefore only possible at the first layer level, where each token can still be unambiguously matched to exactly one channel-window pair of the input signal; this restriction is precisely what allows us to analyze the inner mechanisms of the Foundation Model at the level of individual EEG channels/electrodes, rather than at the level of mixed, uninterpretable intermediate representations.

### 4.1 Limitations

In the current work we focused on non-hardware accelerated pipelines, which discouraged the use of fine-tuning, or the use of the full encoder’s output representations as feature vectors, both of which could plausibly further improve performance at additional computational cost. Literature on the ability of FMs to reconstruct topomaps and region separation [15] has shown that FMs can often be biased by the pretraining dataset or emphasize neighboring connections more. The implications of this has not yet been studied for the REVE model nor for the method applied here. Thus, confirmation of literature on left-right hemisphere asymmetry of positive vs negative stimuli [26] was not possible. Further research should also be conducted on window-based weighted functional connectivity features rather than trial aggregation, as well as other models and additional datasets beyond SEED-IV, to determine the universality of both the symmetry-based head-selection criterion and the frontal-region enrichment finding.

## 5 Conclusion

We have shown that the first-layer attention maps of a pretrained EEG Foundation Model, REVE, can be aggregated over the duration of a trial into channel-level soft functional connectivity matrices. We provide a label-free symmetry criterion, derived from the known tendency of trained encoder-decoder transformers to develop diagonally symmetric attention patterns [19], to identify which attention head carries class discriminative information. On the SEED-IV emotion-recognition benchmark, classical machine learning classifiers trained on this single head’s attention-generated weighted soft functional connectivity matrix are competitive with several fully fine-tuned state-of-the-art EEG Foundation Models, while requiring on the order of 176*×* fewer calculations than standard all-layer attention pooling of the same model. A regional analysis of the resulting discriminative edges further recovers the expected over-representation of frontal connectivity in affective processing, lending additional face validity to the extracted features. Together, these results indicate that lightweight, interpretable linear probing of early-layer attention maps is a promising and computationally accessible alternative to full fine-tuning of EEG Foundation Models, particularly in settings where hardware acceleration is limited. Future work should extend this analysis to additional BCI datasets and tasks, and investigate whether the same symmetry-based criterion generalizes to other experimental settings.

